# CHITOSAN FROM *Portunus Pelagicus* IN THE SYNTHESIS OF REDUCED GOLD NANOPARTICLE AS POTENTIAL CARRIER FOR THE DELIVERY OF ERYTHROPOIETIN

**DOI:** 10.1101/044875

**Authors:** Maria Angelica M. Duque, Rhowell N. Tiozon, Rebecca C. Nueva España

## Abstract

Nanotechnology and its promises for clinical translation to targeted drug delivery with limited accompanying toxicity provide exciting research opportunities that demands multidisciplinary approaches. The colloidal metallic systems have been recently investigated in the area of nanomedicine. Gold nanoparticles have found themselves useful for diagnostics and drug delivery applications. In this study, we have reported a novel method for the synthesis of gold nanoparticles using natural, biocompatible and biodegradable chitosan which came from deacetylating chitin from Portunus Pelagicus. It serves many purposes, as a reducing agent, stabilizer and absorption and penetration enhancer.

Erythropoietin would have high loading efficiency with chitosan reduced gold nanoparticles; the binding is predominantly through hydrogen bonding. Chitosan reduced gold nanoparticles improve the pharmacodynamics and cellular uptake of Erythropoietin across mucosal sites and have immunoadjuvant properties.

There is almost 50 % shell waste in crustacean industry. It is resourceful if it would be bioconverted. The process of bioconversion is deproteination, demineralization and deacetylation to obtain chitosan. In synthesizing gold nanoparticles, 1.48 × 10^−2^ M chloroauric acid will be reduced by heating for 15 minutes in 100mL chitosan solution prepared in 1% acetic acid to yield a ruby-red solution. Erythropoietin would be loaded into it and will undergo 13,000rpm of centrifuge followed by calculating the loading efficiency.

## Introduction

Nanotechnology is really promising in terms of its extensive use. In the last few years, there have been considerable researches in the area of nanoparticle-based drug delivery system that carries different sizes of molecules. According to Merriam Webster, “Nanotechnology is the science of manipulating materials on an atomic or molecular scale especially to build microscopic devices.” Its definition implies that the goal of nanotechnology, specifically the “nano-deliveries” is to revolutionalize the drug delivery where it will reduce or limit the side-effects and allow high efficiency in targeting delivery of drug molecules. Indeed, it is a rapidly thriving science that gives way to many open doors to deal with structures that have at least one dimension of the size of one hundred nanometers or less (Panyala, Mendez, Havel 2009)

Many approaches have been presented because of the potential of nanotechnology, one of which is by using Gold-nanoparticles. According to Mahdihassan (1971,1981) gold was already being used by the Chinese as a medicine in 2500 B.C. In India, Cinnabar Gold which is known to them as “Makaradhwaja” is used as drug primarily for youthful vigour in India. It is a form of ayurvedic medicine to rejuvenate and revitalize the people of India. In the 16^th^ century, Paracelsus created a potion called Aurum Potabile that believed to treat epileptic persons (Daniel et al. 2004). In the 19^th^ century, Michael Faraday discovered through experiment that the reason behind the color of gold solutions was the small size of its particles. It was also used to treat syphilis. When 19^th^ century came, many diseases have been believed to be treated by gold and many approaches were applied to develop it. Gold’s properties make it famous throughout the history. It is a noble metal resistant to oxidation and has interesting electrical, magnetic, optical and physical properties. The pure form of it is non-toxic and non-irritating when ingested. Gold nanoparticles have strong optical absorption, good scattering properties and low or complete lack of toxicity. The properties mentioned create a promising class to the world of nanomedecine. Gold is excellent for bioconjugation with biological. Herein we show a novel method for the synthesis of gold nanoparticles using a biocompatible, biodegradable and natural polymer; chitosan. Many researchers have presented that biologically active substances with amine functions can bind strongly with gold nanoparticles. The process of making chitosan is by deacetylating chitin from crustacean shells. Crustacean processing produces 40% of shell waste because Filipinos are innately resourceful, we can recycle it for nanomedicine. One of the purposes of it is absorption enhancer together with the gold nanoparticle and drug.

In statistics conducted worldwide by the US, Erythropoietin has the biggest sale with $9451 and the demand is still growing. Indeed, anemia is one of the most profuse illnesses in the world. Erythropoietin is used in the number of clinical circumstances to treat anemia due to renal failure, cancer, bone marrow transplantation, AIDS etc. Presently Erythropoietin is administered either as an intravenous or subcutaneous injection at least 3 times a week, depending on the patient’s requirement. Surprisingly, it has 20-30% bioavailability only. The goal of our research is to present a carrier that would improve the pharmacodynamic activity of the drug.

As far as the researches are concern, nationwide and worldwide, no studies yet have been conducted focusing on synthesizing reduced gold nanoparticle using chitosan as novel carrier for delivery of Erythropoietin. Employing to this discussion, this study aimed to synthesize reduced gold nanoparticle using chitosan as potential carrier of erythropoietin against anemia.

This study aims to synthesize reduce gold nanoparticle as potential carrier of artemisinin against malaria. Specifically, this study seeks to answer the following questions:

1. What is the potential of chitosan reduced gold nanoparticle to hold erythropoietin in drug delivery?
2. What is the loading efficiency of erythropoietin on chitosan reduced gold nanoparticle?
3. What is the capability of chitosan in synthesizing gold nanoparticle from chloroauric acid?
4. Is there a significant difference between chitosan concentrations in delivering erythropoietin?

### Nanotechnology

Nanotechnology is a multidisciplinary scientific undertaking that involves creating and utilizing the materials, device or system which is on nanometer scale ^[4]^. It is an advanced scientific technique in the 21stcentury that continually catches the attention of many people involved in medicine. When analyzing the relationship between nanotechnology and biological medicine, nanotechnology and bioavailability and the advances together with the existing problems of bioavailability enhancement, the application of nanotechnological methods for the action mechanism of the drug, it is seen that nanotechnology is one of the fastest developmental, the most potential and the far-reaching high and new technologies in current world, and it enhances the drug delivery ^[5]^.

It covers both current work and concepts that are more advanced where it refers to the projected ability to construct items from the bottom up, using techniques and tools being developed today to make complete, high performance products. ^[6]^ It is pronounced in the sense that the size of the system under study becomes smaller. The materials reduced to the nanoscale can show different properties compared to what they exhibit on a macroscale, enabling unique applications. For instance, opaque substances can become transparent (copper); stable materials can turn combustible (aluminum); insoluble materials may become soluble (gold). A material such as gold, which is chemically inert at normal scales, can serve as a potent chemical catalyst at nanoscales. Much of the fascination with nanotechnology stems from these quantum and surface phenomena that matter exhibits at the nanoscale. ^[7]^

An application of nanotechnology is the existence of Nanoparticles with different sizes that have different biomedical purposes. The potential use of polymeric nanoparticles as drug carriers has led to the development of many different colloidal delivery vehicles. The main advantages of this kind of systems lie in their capacity to cross biological barriers, to protect macromolecules, such as peptides, proteins, oligonucleotides, and genes from degradation in biological media, and to deliver drugs or macromolecules to a target site with following controlled release. In the last few years, several synthetic as well as natural polymers have been examined for pharmaceutical applications. The basic requirement for these polymers to be used in humans or animals is that they have to be degraded into molecules with no toxicity for biological environments. In recent years, biodegradable polymers have attracted attention of researchers to be used as carriers for drug delivery systems. ^[8]^ Moreover, nanoparticles are substantially smaller than eukaryotic and prokaryotic cells where their sizes are comparable to that of antibodies and viruses. It can enter into smallest capillary vessels due to the ultra-tiny volume size they have. It also avoids rapid clearance by phagocytes so that their duration in the bloodstream would be prolonged greatly ^[4]^.

The considerable researches about the nanoparticles-based drug delivery systems involve drugs which are dissolved, entrapped, encapsulated or attached to nanoparticles matrices. Nanoparticle-based drug are believed to alter and improved the pharmacokinetic and pharmacodynamic properties of various type of drug molecules. There are different approaches for drug delivery applications however the remnants of conventional drug-delivery systems are dendrimers, solid-lipid nanoparticles, polymeric nanoparticles, polymeric micelles, liposomes, nanosuspensions and nanocrystals, ceramic nanoparticles, carbon nanotubes (CNTs), quantum dots (QDs), polymersomes and gold nanoparticles ^[4]^.

### Gold nanoparticles

Colloidal gold nanoparticles consist of an elemental gold core surrounded by a negative ionic double layer of charges. Specifically, gold colloidal particles are composed of an internal and essentially crystalline core of pure gold (Au), on the surface of which are adsorbed negative ions, which constitute the inner layer of the ionic double layer. These ions confer the negative charge to the colloidal gold and prevent particle aggregation by electrostatic repulsion. ^[6]^

Gold has long been known to form stable colloids of nanosize particles. Such nanoparticles exhibit different colours due to surface plasmon resonance effects which result from the resonance interaction of incoming electromagnetic radiation in the visible region with the collective plasmon oscillations at the metal surface ^[7]^. Generally, gold nanoparticles are produced in a liquid (“liquid chemical methods”) by reduction of chloroauric acid (H[AuCl4]). After dissolving H[AuCl4], the solution is rapidly stirred while a reducing agent is added. This causes Au3+ ions to be reduced to neutral gold atoms. As more and more of these gold atoms form, the solution becomes supersaturated, and gold gradually starts to precipitate in the form of sub-nanometer particles. The rest of the gold atoms that form stick to the existing particles, and, if the solution is stirred vigorously enough, the particles will be fairly uniform in size. To prevent the particles from aggregating, some sort of stabilizing agent that sticks to the nanoparticle surface is usually added. Alternatively, gold colloids can be synthesised without stabilizers by laser ablation in liquids ^[8]^.

Colloidal gold suspensions consist of small granules of this transition metal in a stable, uniform and dispersion viewed under the electron microscope they appear as solid spheres of dense material. In light microscopy meanwhile, the particles appear as light dots on a darker background due to the high reflections of the particles or as an orange red coating where they are localize in large conglomerates in cell or tissues.^[6] [22] [25]^ Colloidal gold preparation

Varieties of different methods have been introduced for the synthesis of colloidal gold. In all the methods developed to date, the emphasis has been on the production of gold colloids that have uniform and controllable diameter in a simple approach. The methods have also produced in nano dispersed gold particles in the 3.30nm range. These methods also share the use of tetrachloroauric acid (HAuCl_4_) but vary considerably with respect to reducing agents, order of reagent addition, physical parameters (concentration, temperature and mixing rate) and of course the resulting diameter of colloidal gold particles.

There are several considerations to take into account in preparing colloidal gold are based on controlled reduction of an aqueous solution of tetrachloroauric acid, then using different reducing agents under varying conditions would yield different colloidal gold sizes. To create large particle colloidal gold dispersions, chloroauric acid normally is treated with sodium citrate. The result is particle range of about 15-150 nm, depending on the concentration of citrate utilized. However in this research, chitosan would be used as reducing agent. ^[9]^

The final diameter of the colloidal gold is determined by the number of nuclei formed at the beginning of the reaction compared with the subsequent rate of shell condensations. The use of rapid reductants (white phosphorus or tannic acid) results in a greater number of nuclei formed, thereby consuming much of the HAuCl_4_ and limiting the amount remaining for shell growth through condensation.

Since colloidal particles prepared in the solution phase have a tendency to agglomerate, it is necessary to protect them using surfactants or polymeric ligands such as polyvinylpyrrolidone (PVP). Varieties of reducing agents have been employed for reductions. These include electrides, alcohols, glycols, and certain specialized reagents such as tetrakis (hydroxyl methyl) phosphonium chloride. The last reagent has enabled the preparation of gold hydrosols (down to ~2nm. ^[26][27]^

### Gold Nanoparticle Probes in Medical diagnostics

The inherent characteristics and properties of colloidal gold probes render them ideal in the field of medicine. Aside from the ease of preparation, prompt bioconjugation with certain macromolecules and cost-effectiveness, the unique optical properties of colloidal gold and the stability they provide for antibodies, enzymes or nucleic acids make them exceptional candidates for use in medical diagnostics. The colour of nanoparticles varies with their size and shapes. Gold atoms can aggregate themselves under various conditions and can form GNPs by sequential aggregation. Gold clusters can be formed by various methodologies. For example, charged gold (Aun+) clusters are formed by laser desorption/ionization (LDI) of Au(s) or Au salts such as auric acid (HAuCl4) where Aun + clusters with n up to 1–25 were detected (Peña-Méndez et al. 2008). An example of mass spectra concerning the formation of gold clusters is given in Figure 1.

**Figure 1.**
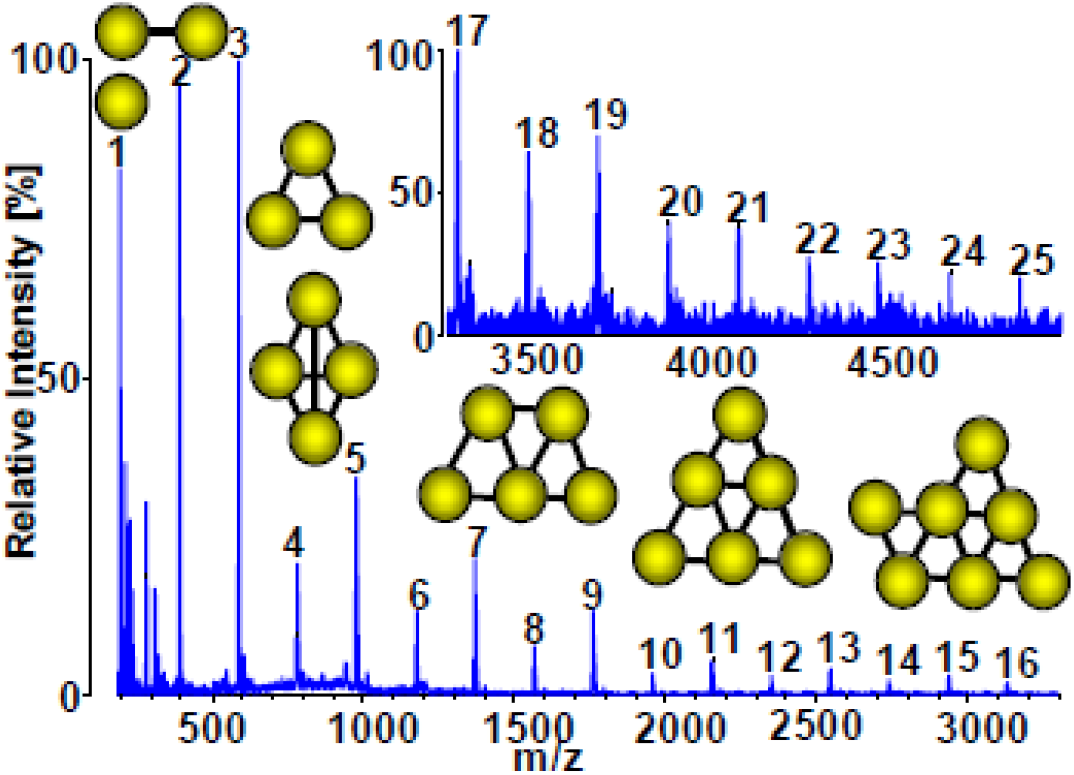
Part of Laser Desorption Ionisatioin TOF mass spectrum showing the formation of gold clusters Au_n_^+^ (n=1-25) ^[11]^.

The spread of numerous diseases, such as cancer and HIV have lead to the explosion of the studies that seek to provide immediate medical diagnostics. Studies in immunocytochemistry and cellular imaging have been constantly sought for improvement and potential application in cancer diagnostics.^[1][6]^Gold is used because its core is essentially inert and non-toxic and exploit unique physical and chemical properties for transporting and unloading pharmaceuticals.^[4]^ The administration of hydrophobic drugs require molecular encapsulation and it is found that nanosized particles are particularly efficient in evading the reticuloendothelial system ^[11][28][29]^

Gold nanoparticles have the ability to resonantly scatter visible and near infrared light upon the excitation of their surface Plasmon oscillation. This properly enables the utilization of gold particles as promising markers in cellular imaging. The scattering light intensity is extremely sensitive to the size and aggregation state of the particles. Preliminary studies have reported their use as constant agents for biomedical imaging using different microscopy techniques such as confocal scanning electron microscopy, multiphoton Plasmon resonance microscopy and optical coherence microscopy. ^[10]^

### Gold nanoparticles in delivery

Gold nanoparticles (AuNPs) provide non-toxic carriers for drug and gene delivery applications. With these systems, the gold core imparts stability to the assembly, while the monolayer allows tuning of surface properties such as charge and hydrophobicity. An additional attractive feature of AuNPs is their interaction with thiols, providing an effective and selective means of controlled intracellular release.

Nanocarriers have provided a novel platform for target-specific delivery of therapeutic agents. Over the past decade, several delivery vehicles have been designed based on different nanomaterials, such as polymers, dendrimers, liposomes, nanotubes, and nanorods. Gold nanoparticles (GNPs) have recently emerged as an attractive candidate for delivery of various payloads into their targets. The payloads could be small drug molecules or large biomolecules, like proteins, DNA, or RNA (vide post). Efficient release of these therapeutic agents is a prerequisite for effective therapy. The release could be triggered by internal (e.g. glutathione(GSH), or pH or external (e.g. light) stimuli. Significantly the internal stimuli operate in a biologically control manner, whereas the external stimuli provide spatio-temporal control over the release. Gold nanoparticles exploit their unique chemical and physical properties for transporting and unloading the pharmaceuticals. First, the gold core is essentially inert and non-toxic. A second advantage is their ease of synthesis; monodisperse nanoparticles can be formed with core sizes ranging from 1 nm to 150 nm.^[33][34][37][39]^

Further versatility is imparted by their ready functionalization, generally through thiol linkages (vide post). Moreover, their photophysical properties could trigger drug release at remote place. Several reviews describing nanoparticle–biomacromolecule interactions have recently been published, generally focusing on biosensing and diagnostic applications. This review will focus on drug, gene, and protein delivery using GNPs.

Gold nanoparticles are capable of delivering large biomolecules, without restricting themselves as carriers of only small molecular drugs. Tunable size and functionality make them a useful scaffold for efficient recognition and delivery of biomolecules. They have shown the success in delivery of peptides, proteins, or nucleic acids like DNA or RNA. Gene therapy presents an ideal strategy for the treatment of genetic as well as acquired diseases. Viruses provide a logical vehicle for gene therapy, and have been shown to be highly efficient. Viruses have nonetheless raised many safety concerns arising from unpredictable cytotoxicity and immune responses.

Synthetic DNA delivery systems have overcome these limitations. However, current non-viral gene delivery systems suffer from less efficiency. The vectors encounter numerous barriers between the site of administration and localization in the cell nucleus. The effective delivery vehicles need to provide efficient protection of nucleic acid from degradation by nucleases, efficient cell entry, and release of the nucleic acid in functional form in the nucleus. (Partha Ghosh et.al, 2008, Gold nanoparticles in delivery application pp.1307-1312) ^[22][25][26][27]^

**Figure 2.**
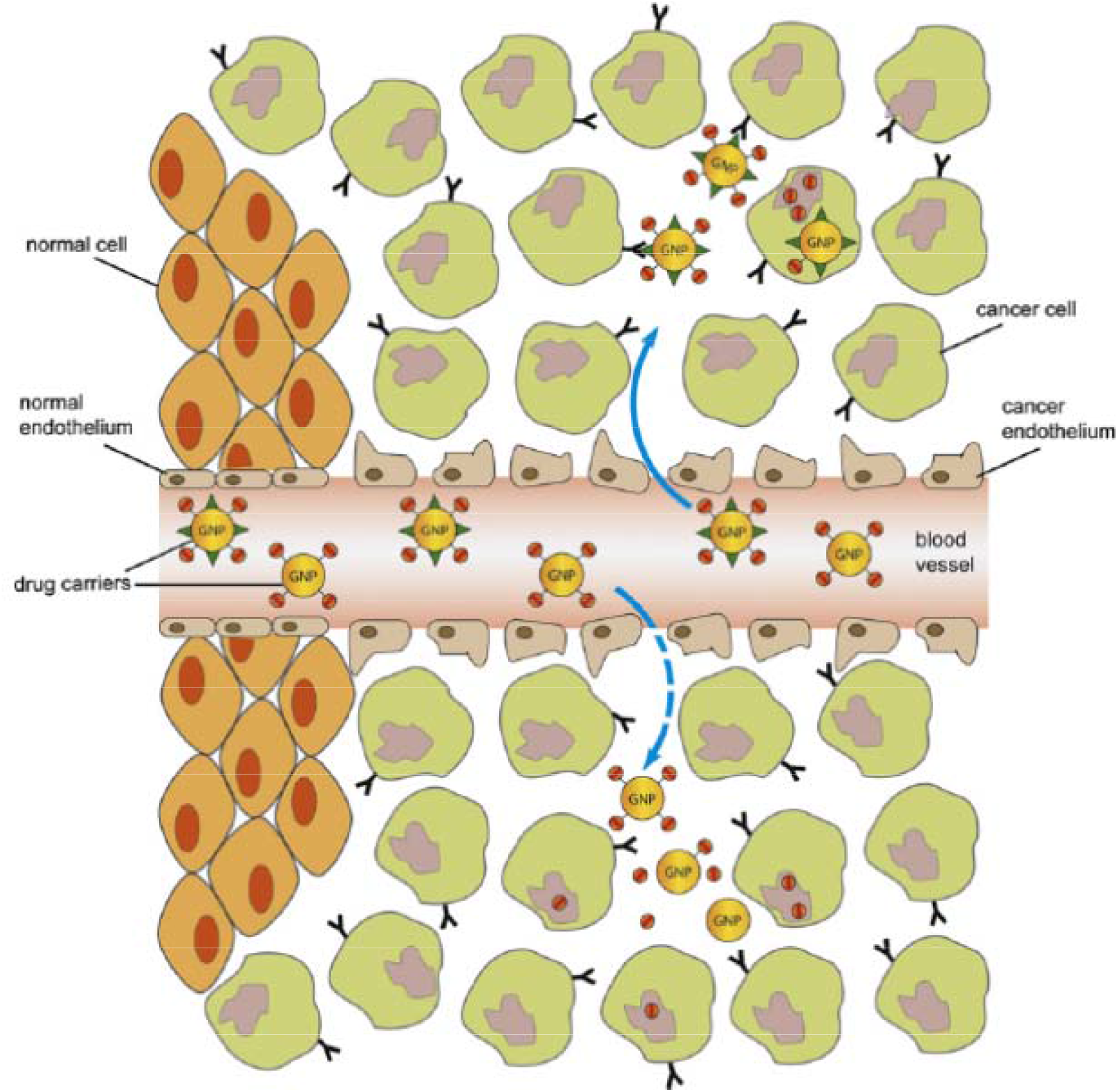
Illustration of drug delivery via “active” and “passive” targeting, solid and dotted line respectively This gives us the picture of the fate of gold nanoparticles and its way of delivering drugs.

### Surface Plasmon Resonance

The SPR is a quantum phenomenon. It is a coherent excitation of all the “free electrons” within the conduction band leading to an in-phase oscillation. When the size of a metal nanoparticle is smaller than the wavelength of incident radiation, a SPR is generated.^[31][32]^The SPR depends heavily on the various intrinsic and environmental factors of the particles such as composition, size, shape, aggregation state, and dielectric medium. This therefore allows for optical tunability of the metal nanoparticle over a range of wavelengths. ^[6]^

### IR Spectroscopy

Infrared spectroscopy is certainly one of the most important analytical techniques available to today’s scientists. One of the great advantages of infrared spectroscopy is that virtually any sample in virtually any state may be studied. Liquids, solutions, pastes, powders, films, fibres, gases and surfaces can all be examined with a judicious choice of sampling technique. As a consequence of the improved instrumentation, a variety of new sensitive techniques have now been developed in order to examine formerly intractable samples.

The infrared portion of the electromagnetic spectrum is usually divided into three regions; the near-, mid- and far- infrared, named for their relation to the visible spectrum. The higher-energy near-IR, approximately 14000–4000 cm-1 (0.8–2.5 μm wavelength) can excite overtone or harmonic vibrations. The mid-infrared, approximately 4000–400 cm-1 (2.5–25 μm) may be used to study the fundamental vibrations and associated rotational-vibrational structure. The far-infrared, approximately 400–10 cm-1 (25–1000 μm), lying adjacent to the microwave region, has low energy and may be used for rotational spectroscopy. The names and classifications of these subregions are conventions, and are only loosely based on the relative molecular or electromagnetic properties. ^[40]^

### Characteristics of Chitin and Chitosan

Chitin is the second most ubiquitous natural polysaccharide after cellulose on earth and is composed of β(1,4)-linked 2-acetamido-2-deoxy-β-D-glucose ^[15]^ (N-acetylglucosamine) (Figure 1). It is often considered as cellulose derivative, even though it does not occur in organisms producing cellulose. It is structurally identical to cellulose, but it has acetamide groups (-NHCOCH_3_) at the C-2 positions. Chitin is a white, hard inelastic, nitrogenous polysaccharide found in the exoskeleton as well as in the internal structure of invertebrates.

**Figure 3.**
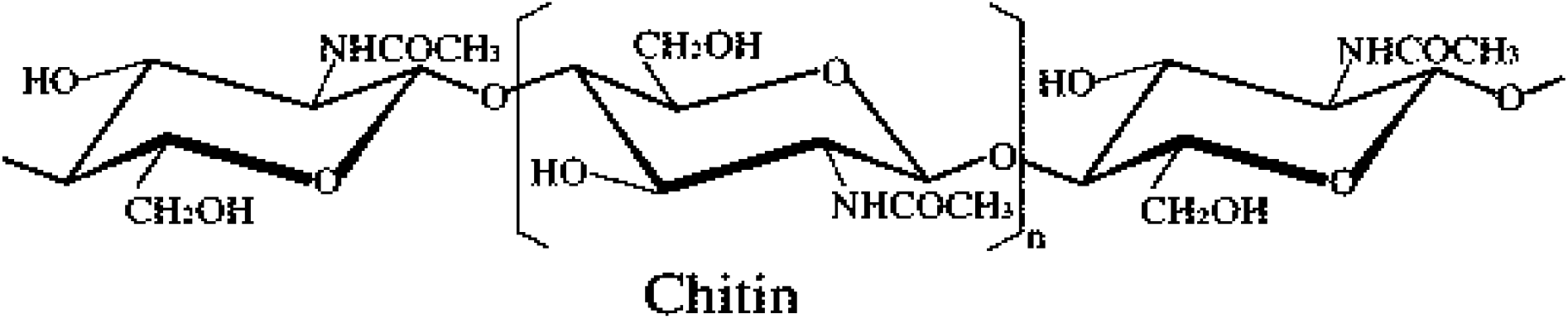
Structure of Chitin

Similarly the principle derivative of chitin, chitosan is a linear polymer of α (1,4)-linked 2-amino-2-deoxy-β-D-glucopyranose and its easily derived by N-deacetylation, to a varying extent that is characterized by the degree of deacetylation (DD), and is consequently copolymer of N-acetylglucosamine and glucosamine (Figure 2). It is one of the most used polysaccharides in the design of drug delivery strategies for administration of either biomacromolecules or low molecular weight drugs. Chitosan and its derivatives in the last two decades have proven to be excellent and safe candidates for improving mucosal and trans-mucosal delivery or drugs, mainly due to their mucoadhesive and absorption enhancing properties, closely related with the cationic character of the polymer ^[15]^. Chitosan has been indicated as a promising biomaterial for biomedical and pharmaceutical, i.e., drug delivery, applications. The characterization of its biocompatibility pattern encompasses, therefore, a major issue, as it will drive the process of a future human drug delivery application. ^[16]^

**Figure 4.**
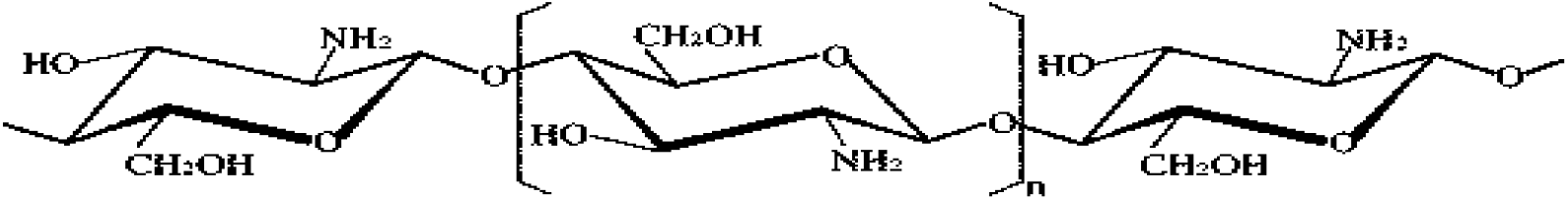
Chitosan

### Properties of Chitin and Chitosan

Most of the naturally occurring polysaccharides e.g, cellulose, dextrin, pectin, alginic acid, agar, and carragenas are naturally and acidic in nature, whereas chitin and chitosan are examples of highly basic polysaccharides. Their properties include solubility in various media, solution, viscosity, polyelectrolyte behavior, polyxysalt formation, ability to form films, metal chelations, optical and structural characteristics ^[17][31]^.

Although the β(1,4)-anhydroglucosidic bond of chitin is also present in cellulose the characteristic properties of chitin/chitosan are not shared by cellulose. ^[17]^ Chitin is highly hydrophobic and is insoluble in water and most organic solvents. It is soluble in hexafluoroisopropanol, hexafluoroacetone, and chloroalcohols in conjunction with aqueous solutions of mineral acids^[18]^. Depending on the extent of deacetylation, chitin contains 5 to 8 percent (w/v) nitrogen, which is mostly in the form of primary aliphatic amino groups as found in chitosan. Chitosan undergoes the reactions of amines, of which N-acetylation and Schiff reactions are most important. Chitosan glucans are easily obtained under mild conditions but it is difficult to obtain cellulose glucans.

Chitosan from simple aldehydes produce N-alkyl chitosan upon hydrogenation. The presence of the more or less bulky substituent weakens the hydrogen bonds of chitosan; therefore, N-alkyl chitosans swell in water inspite of the hydrophobicity ofalkyl chains. They retain the film forming property of chitosan^[15]^.

### Chitin Extraction

As mentioned above, chitin is present within numerous taxonomic groups. However, commercial chitins are usually isolated from marine crustaceans, mainly because a large amount of waste is available as a by-product of food processing. In this case, α-chitin is produced while squid pens are used to produce β-chitin. The structure of α-chitin has been investigated more extensively than that of either the β- or the γ- form, because it is the most common polymorphic form. Very few studies have been carried out on γ- chitin. It has been suggested that γ-chitin may be a distorted version of either α- or β-chitin rather than a true third polymorphic form ^[12]^

In α -chitin, the chains are arranged in sheets or stacks, the chains in any one sheet having the same direction or ’sense’. In β-chitin, adjacent sheets along the *c* axis have the same direction; the sheets are parallel, while in α-chitin adjacent sheets along the *c* axis have the opposite direction, they are antiparallel. In γ- chitin, every third sheet has the opposite direction to the two preceding sheets ^[13]^.

### Chitin Deacetylation

Chitosan is prepared by hydrolysis of acetamide groups of chitin. This is normally conducted by severe alkaline hydrolysis treatment due to the resistance of such groups imposed by the trans arrangement of the C2-C3 substituents in the sugar ring ^[13]^. Thermal treatments of chitin under strong aqueous alkali are usually needed to give partially deacetylated chitin (DA lower than 30%), regarded as chitosan. Usually, sodium or potassium hydroxides are used at a concentration of 30-50% w/v at high temperature (100ºC). In general, two major different methods of preparing chitosan from chitin with varying degree of acetylation are known. These are the heterogeneous deacetylation of solid chitin and the homogeneous deacetylation of pre-swollen chitin under vacuum (by reducing pressure) in an aqueous medium.

**Figure 5.**
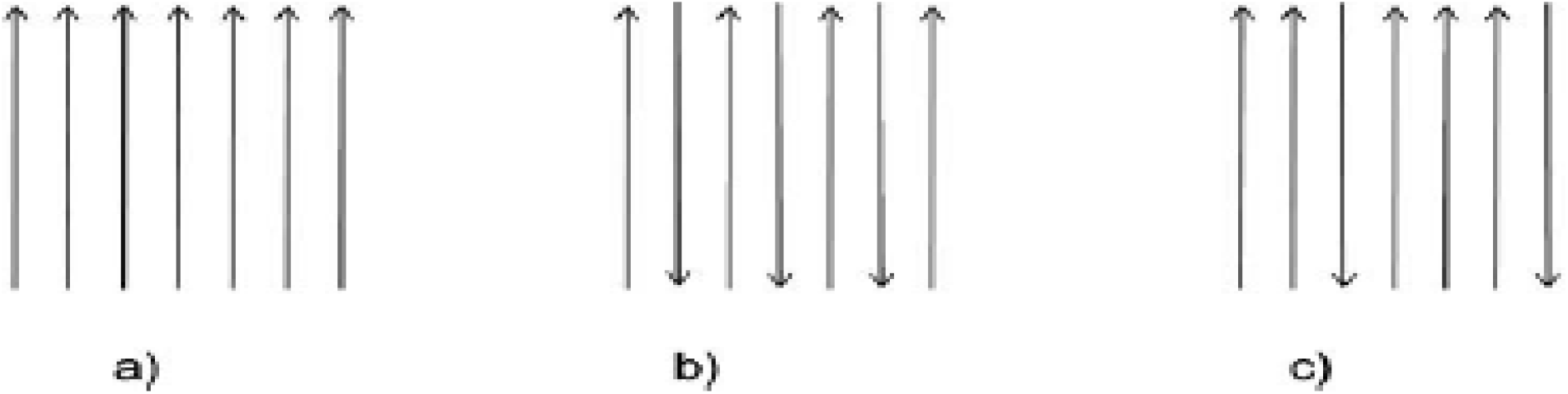
Three polymorphic configurations of Chitin (A) α-chitin, (B) ß-chitin and (C) γ-Chitin ^[13]^

In both, heterogeneous or homogeneous conditions, the deacetylation reaction involves the use of concentrated alkali solutions and long processing times which can vary depending on the heterogeneous or homogeneous conditions from 1 to nearly 80 hours. Factors that affect the extent of deacetylation include concentration of the alkali, previous treatment, particle size and density of chitin. The last two factors affect penetration rate of the alkali into the amorphous region and to some extent also into the crystalline regions of the polymer, needed for the hydrolysis to take place. In practice, the maximal DD that can be achieved in a single alkaline treatment is about 75-85% ^[13]^. In general, during deacetylation, conditions must be the proper ones to deacetylate, in a reasonable time, the chitin to yield a chitosan soluble in diluted acetic acid.

### Conversion of chitin to chitosan

A schematic representation of the processes to prepare chitin and chitosan from raw material is shown in Scheme 1.

**Scheme 1.**
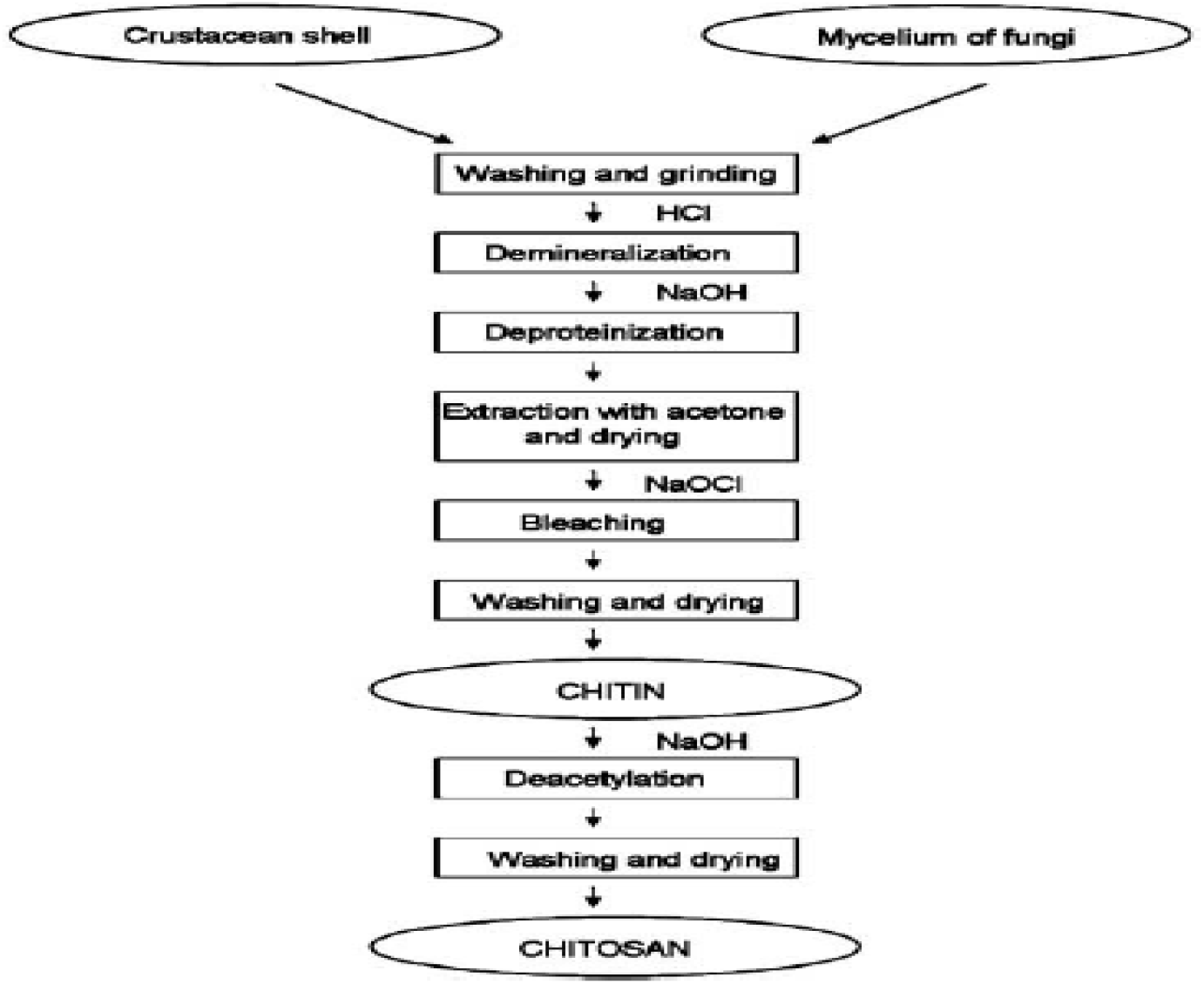
Preparation of chitin and chitosan from raw material.

Crustacean shells consist of 30-40% proteins, 30-50% calcium carbonate, and 20-30% chitin and also contain pigments of a lipidic nature such as carotenoids (astaxanthin, astathin, canthaxanthin, lutein and β-carotene). These proportions vary with species and with season. On the other hand, β-chitin is associated with a higher protein content but lower carbonate concentration. Chitin is extracted by acid treatment to dissolve the calcium carbonate followed by alkaline extraction to dissolve the proteins and by a depigmentation step to obtain a colourless product mainly by removing the astaxantine. ^[14]^

### Biological properties of chitin and chitosan

Chitin and chitosan are currently receiving a great deal of interest as regards medical and pharmaceutical applications because they have interesting properties that make them suitable for use in the biomedical field, such as biocompatibility, biodegradability and non toxicity. Moreover, other properties such as analgesic, antitumor, hemostatic, hypocholesterolemic, antimicrobian, and antioxidant properties have also been reported. ^[13]^ A deeperunderstanding of the mechanism of these properties makes it necessary for chitosan to be well characterized and purified from accompanying compounds. ^[14]^ In addition, chitins and chitosans derivatized in a variety of fashions can be used to prove molecular hypothesis for the biological activity. Since the majority of the biological properties are related to the cationic behaviour of chitosan, the parameter with a higher effect is the DD. However, in some cases, the Mw has a predominant role. ^[31]^

### Applications of Chitin and Chitosan

The chitin originates from the study of the behavior and chemical characteristics of lysozome, an enzyme present in the human body fluids. It dissolves certain bacteria by cleaving the chitinous material of the cell walls. ^[20]^ A wide variety of medical applications for chitin and chitin derivatives have been reported over the last three decades. It has been suggested that chitosan may used to inhibit fibroplasias in wound healing and to promote tissue growth and differentiation in tissue culture. ^[21]^

### Biomedical Applications of Chitosan

The design of artificial kidney systems has made possible repetitive hemodialysis and the sustaining life of chronickidney failure patients. Chitosan membranes have been proposed as an artificial kidney membrane because of their suitable permeability and high tensile strength^[22]^. The most important part of artificial kidney is the semipermeable membrane. These novel membranes need to be developed for better control of transport, ease of formability and inherent blood compatibility.

### Drug Delivery Systems

An important application of chitosans in industry is the development of drug delivery systems such as nanoparticles, hydrogels, microspheres, films and tablets (Figure 3). As a result of its cationic character, chitosan is able to react with polyanions giving rise to polyelectrolyte complexes ^[23]^. Pharmaceutical applications include nasal, ocular, oral, parenteral and transdermal drug delivery. Three main characteristics of chitosan to be considered are: Molecular weight, degree of acetylation and purity. When chitosan chains become shorter (low Mw chitosan), it can be dissolved directly in water, which is particularly useful for specific applications in biomedical or cosmetic fields, when pH should stay at around 7.0.

In drug delivery, the selection of an ideal type of chitosan with certain characteristics is useful for developing sustained drug delivery systems, prolonging the duration of drug activity, improving therapeutic efficiency and reducing side effects. It is suggested that the physicochemical characteristics of chitosan are important for the selection of the appropriate chitosan as a material for drug delivery vehicles ^[21]^. Investigations have indicated that DD and Mw of chitosan have significantly affected the role of chitosan in therapeutic and intelligent drug delivery systems ^[20][28][30][36]^.

**Figure 7.**
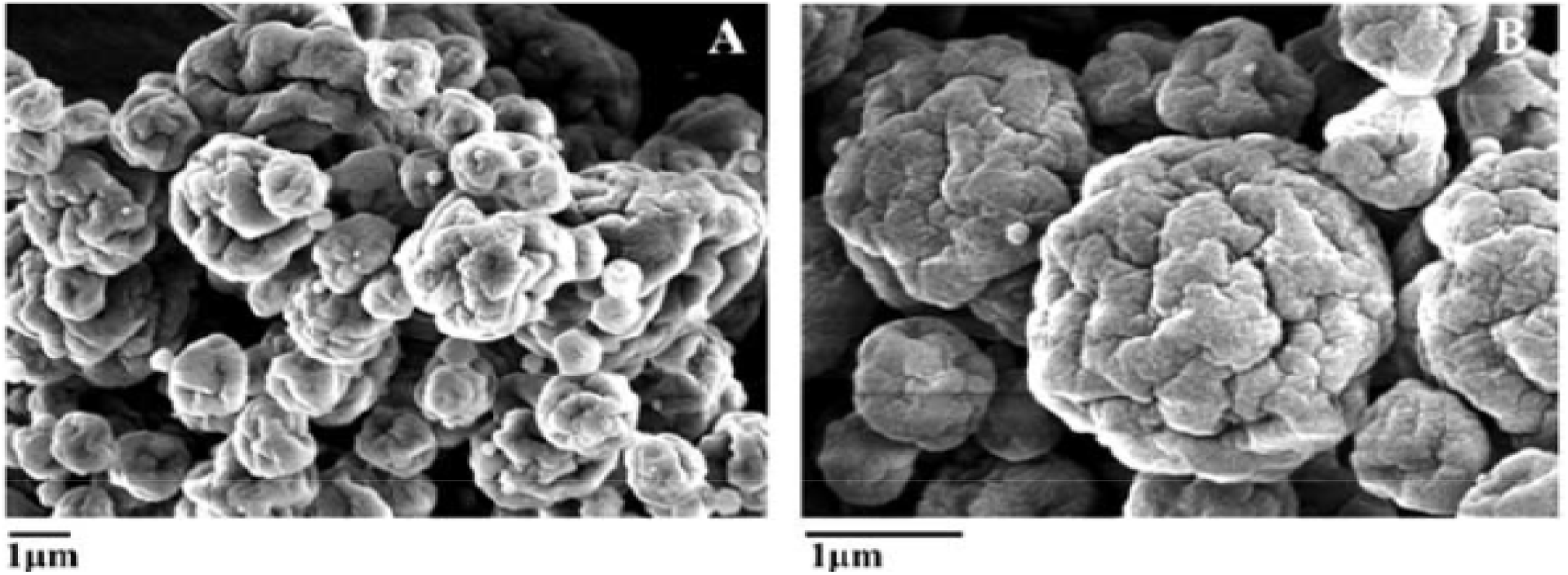
(A) High Mw Chitosan (640 kDa) microspheres crosslinked with 0.2% TPP obtained by spray-drying. (B) Detail of the microspheres ^[15]^.

The DD controls the degree of crystallinity and hydrophobicity in chitosan due to variations in hydrophobic interactions which control the loading and release characteristics of chitosan matrices. The DD also controls the degree of cross-linking of chitosan in the presence of any suitable cross-linker. The higher the DD is, the higher the number of free amino groups and therefore the degree of covalent cross-linking increases ^[16]^. When analyzing the influence of cross-linking degree and degree of deacetylation on size and morphology of the microspheres, these authors reported that the size and the surface roughness decreased on increasing the degree of cross-linking and the degree of deacetylation. It is also reported that a high degree of chitosan deacetylation and narrow polymer Mw distribution were shown to be critical for the control of particle size distribution ^[17][32][33][34][37]^.

A higher degree of cross-linking and a higher DD in chitosan increase the compactness of matrices and its hydrophobicity, thus controlling the degree of swelling and diffusivity of the drug entrapped in chitosan matrixes. It was observed that a DD between 48-62% promotes maximal loading capacity, due to the size of the cross-links and pores formed. Regarding the release properties, a very low DD can induce burst release ^[21]^.

The microspheres with medium Mw chitosan showed an optimum loading efficiency ^[19]^. Microsphereswith medium Mw chitosan are more efficient in releasing the centchroman in a controlled manner in comparison to low and high Mw chitosan microspheres. The initial burst release of centchroman from microspheres with different Mws and different degrees of deacetylation of chitosan varied linearly with the square root of the release time indicating a diffusion-controlled release of centchroman from these microspheres (n = 0.5). The influence of chitosan DD and Mw on the microspheres properties prepared as matrix for drug delivery is shown in Table 1.

**Table 1.** Influence of Chitosan DD and Mw on Microspheres Properties

| Physico-Chemical Property | Effect on Microsphere Properties |
| --- | --- |
| □ DD | ↑ covalent crosslinking<br>↓ size<br>↓ surface roughness<br>↓ swelling<br>↑ compactness and hydrophobicity<br>↓ loading capacity<br>↓ burst release |
| □ Mw | ↑ sphericity<br>↑ morphology homogeneity<br>↑ crosslinking<br>↓ swelling<br>↓ release rate<br>↓ diffusion coefficient (D) |

The previous researchers studied the relationship between physicochemical characteristics and functional properties of chitosan such as the ability to form spherical gel, control of drug release from chitosan gel and biodegradation of chitosan ^[20]^. They found that the formation of spherical chitosan gels in aqueous amino acid solution or aqueous solution containing metal ions was affected mainly by viscosity of the chitosan solution. High concentration of chitosan species with a high Mw could not be used to prepare chitosan spherical gel due to its high viscosity and the use of very low concentration of chitosan did not result in instantaneous spherical gel formation because the diffusion of chitosan within the preparative medium was too rapid. The degree of deacetylation also had an effect on spherical gel formation in the case of gelation of chitosan by chelation with metal ions. Chitosan with high degree of deacetylation was able to form spherical gel by chelation due to higher availability of amino groups that chelated with metal ions better than chitosan of low DD. Only in the case of chelation with metal ions was the extent of deacetylation related to drug release. ^[30][35][36]^

### Portunus Pelagicus

The famous type of crab here in the Philippines is called “Alimasag”. It is also known as also known as the flower crab, blue crab, blue swimmer crab, blue manna crab or sand crab. The name “flower crab” is used in east Asian countries while the latter names are used in Australia. The crabs are widely distributed in eastern Africa, Southeast Asia, East Asia, Australia and New Zealand.

They stay buried under sand or mud most of the time, particularly during the daytime and winter, which may explain their high tolerance to ammonium (NH4+) and ammonia (NH3). They come out to feed during high tide on various organisms such as bivalves, fish and, to a lesser extent, macroalgae. They are excellent swimmers, largely due to a pair of flattened legs that resemble paddles. However, in contrast to another portunid crab (Scylla serrata), they cannot survive for long periods out of the water. The species is commercially important throughout the Indo-Pacific where they may be sold as traditional hard shells, or as “soft-shelled” crabs, which are considered a delicacy throughout Asia. The species is highly prized as the meat is almost as sweet as Callinectes sapidus.

These characteristics, along with their fast growth, ease of larviculture, high fecundity and relatively high tolerance to both nitrate and ammonia, (particularly ammoniacal nitrogen, NH3–N, which is typically more toxic than ammonium, as it can more easily diffuse across the gill membranes), makes this species ideal for aquaculture. ^[24]^

### Erythropoietin

Erythropoietin (EPO) is a glycoprotein hormone produced primarily by the kidney in response to hypoxia and is the key regulator of red blood cell (RBC) production. EPO is involved in all phases of erythroid development, and has its principal effect at the level of erythroid precursors. After EPO binds to its cell surface receptor, it activates signal transduction pathways that interfere with apoptosis and stimulates erythoid cell proliferation. Erythropoietin stimulates erythropoiesis in anaemic patients with chronic renal failure in whom the endogenous production of erythropoietin is impaired. Because of the length of time required for erythropoiesis – several days for erythroid progenitors to mature and be released into the circulation – a clinically significant increase in haemoglobin is usually not observed in less than two weeks and may require up to ten weeks in some patients. (Eprex, Jansenn et. al 2012, Subcutaneous and Intravenous injection: Product information)

Since the innovation of recombinant DNA technology, the pharmaceutical industry is capable of producing highly purified therapeutic protein and peptide drugs. However, the major route of administration is still parenteral route. With the use of recombinant DNA technology, recombinant human EPO could be manufactured in large quantities for therapeutic purposes. Recombinant human EPO has a molecular weight of 30kDa, activity due to those of natural EPO isolated from human urine. Studies are carried out through different route like oral, nasal and rectal among those, oral is the most attractive. (Naranjan Ventakasan et. Al 2005, Liquid fill nanoparticles as a drug delivery tool for protein therapeutic) ^[23][28][36][38]^

### Chitosan reduced gold nanoparticles as novel carrier

This research was made by Bhumkar, Joshi, Sastry and Porkharkar. They used chitosan as reducing agent in synthesising gold nanoparticles from chloroauric acid. The drug used here is insulin which was loaded onto chitosan. Chitosan is known to act as a penetration enhancer for proteins and vaccines administered across the mucosal routes. It was used because of its compatibility, chemical nature and absorption enhancement properties. The reason chitosan can be a reducing agent is its electronegativity property. It would act as electrostatic stabilizer because it is a polyelectrolyte. Having said that, it would render dual advantage by providing sufficient charge through amino groups which will aid in the subsequent attachment of the biomolecules as well render optimum stability and subsequently help to improve the uptake of the nanoparticles. Gold has been used because it shows different optical properties at nanosclae than its bulk counterpart. These properties are dependent on composition, size, shape and surrounding medium of the particles. The surface Plasmon band of gold nanoparticles comes in visible region that’s why it can be used to monitor shape, size and aggregation of nanoparticles. The dipole resonance from the neighbouring nanoparticles overlap and it makes the surface Plasmon band sensitive to interparticle distance. In order to elucidate the cause of aggregation at lower chitosan concentrations, zeta potential measurements were carried out. Zeta potential involves the charge of gold nanoparticles before and after loading the drug. The measurements of zeta potential allow predictions about the stability of colloidal aqueous dispersions. Usually, particle aggregation is less likely to occur for charged particles with optimum zeta potential (” > 30 mV) due to electrostatic repulsions. The zeta potential and standard deviation values of gold nanoparticles before and after loading of insulin are shown in the table.

**Table 2.** Zeta Potential of Blank Gold Nanoparticles Synthesized Using Varying Concentrations of Chitosan and Insulin Loaded Chitosan Reduced Gold Nanoparticles

| Gold Nanoparticles Reduced with Different Chitosan Concentrations | Zeta Potential (mV) |
| --- | --- |
| 0.01% | +4.23 |
| 0.05% | +7.02 |
| 0.1% | +58.3 |
| 0.2% | +62.7 |
| 0.5% | +51.3 |
| 1% | +50.2 |
| Insulin loaded gold nanoparticles reduced with 0.2% chitosan | +65.2 |

The nanoparticles synthesized using higher concentrations of chitosan (0.1% w/v– 1% w/v) had no shift in SPB, and showed no aggregation, indicating that nanoparticles formed were stable. Thus, the long-term stability at higher concentration could be attributed to the electrostatic and mechanical barrier properties of chitosan. The viscosity of gold nanoparticles reduced using varying concentrations of chitosan was determined. It was observed that the viscosity was higher at 0.5% w/v chitosan (Table III). Taking into consideration the stability and viscosity of the gold nanoparticles, nanoparticles synthesized using 0.2% w/v chitosan were chosen for subsequent loading of insulin.

The particle size range of the nanoparticles was between 10–50 nm as shown in Figure below. No significant effect on the particle size distribution of gold nanoparticles was observed with varying concentration of chitosan. Insulin loading on the nanoparticles also did not lead to any effect on the particle size. The particle size determined using photon correlation spectroscopy revealed a monodisperse nature of the 0.2% w/v chitosan reduced gold nanoparticles. The hydrodynamic diameter of 0.2% chitosan reduced gold nanoparticles was found to be 100 nm. The loading efficiency of insulin on chitosan reduced gold nanoparticles was found to be 53%. The nature of binding of insulin to the gold nanoparticles plays vital role in the release and subsequent activity. Reduction of chloroauric acid with chitosan results in formation of chitosan coated gold nanoparticles. It was also observed that the insulin loaded nanoparticles showed 4-5 times greater permeation as compared to free insulin. The nanoparticles have the ability to make the loaded protein more stable and protect it from the harsh environment of the gastrointestinal tract. Thus, chitosan reduced gold nanoparticles have resulted in a significant improvement in the uptake.

## Materials and Methods

The materials used in this study were the following reagents: Hydrochloric acid (HCl), Sodium hydroxide (NaOH), Chloroauric acid (HAuCl_4_), acetic acid (CH_3_COOH). The source of chitin was *Portunus Pelagicus* or commonly known as “Alimasag”. The drug that was delivered is erythropoietin.

**Figure 3.1.**
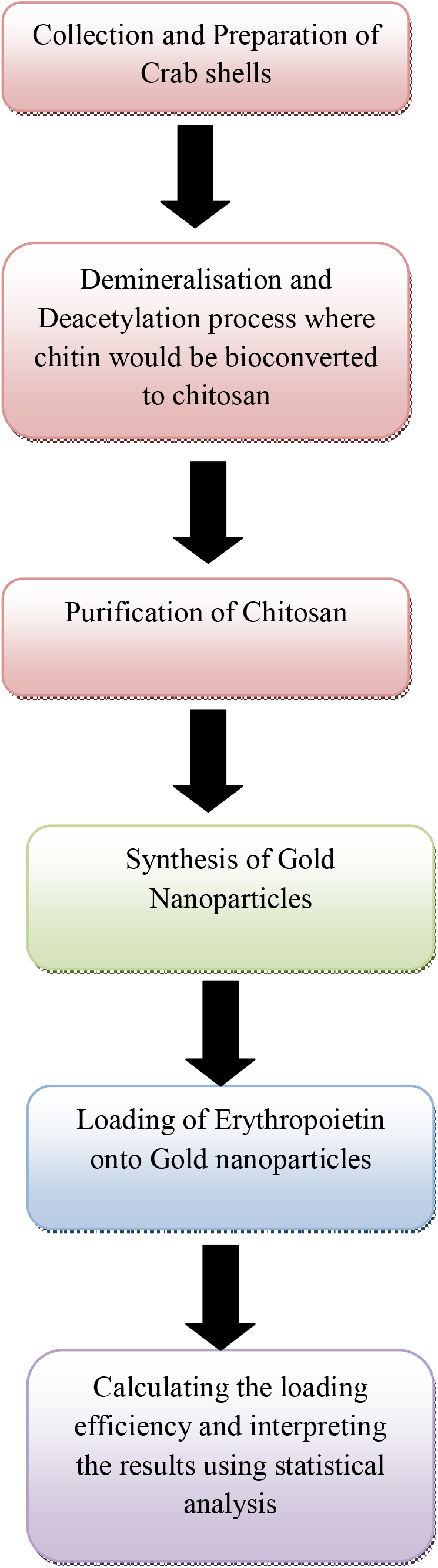
The Schematic Diagram of this

### Collection and Preparation of Crabshells

The crabshells was collected from seafood restaurants. It was washed to remove any excess crab flesh or anything that was not needed in the study. It was authenticated by a Zoologist at National Museum to identify the type of the species. The crab shells were collected in big plastic bags with addition of ice to preserve the crab shell from decomposition. During transportation to the laboratory (transport time was only one and a half hour), the crab waste was kept cool and in the dark to avoid effects of direct sunlight. After arrival, the shrimp waste was directly used to produce chitosan as specified below. In this paper crab shell without further treatment is named fresh crab shell, crab shell stored in the freezer (at minus 180C) for several months is named frozen crab shell.

### Demineralisation and Deacetylation process

The grounded exoskeleton is demineralised using 1% HCl with four times its quantity. The samples were allowed to soak for 24 h to remove the miner-als (mainly calcium carbonate) (Trung et al., 2006). The demineralized shrimp shell samples were then treated for one hour with 50 ml of a 2% NaOH solution to decompose the albumen into water soluble amino-acids. The remaining chitin is washed with deionized water, which is then drained off. The chitin was further converted into chitosan by the process of deacetylation (Huang et al., 2004).

The deacetylation process is carried out by adding 50% NaOH and then boiled at 100°C for 2 h on a hot plate. The samples are then placed under the hood and cooled for 30 min at room temperature. After-wards the samples are washed continuously with the 50% NaOH and filtered in order to retain the solid matter, which is the chitosan. The samples were then left uncovered and oven dried at 110°C for 6 h. The chitosan obtained will be in a creamy-white form (Muzzarelli and Rochetti, 1985).

**Figure 3.2.**
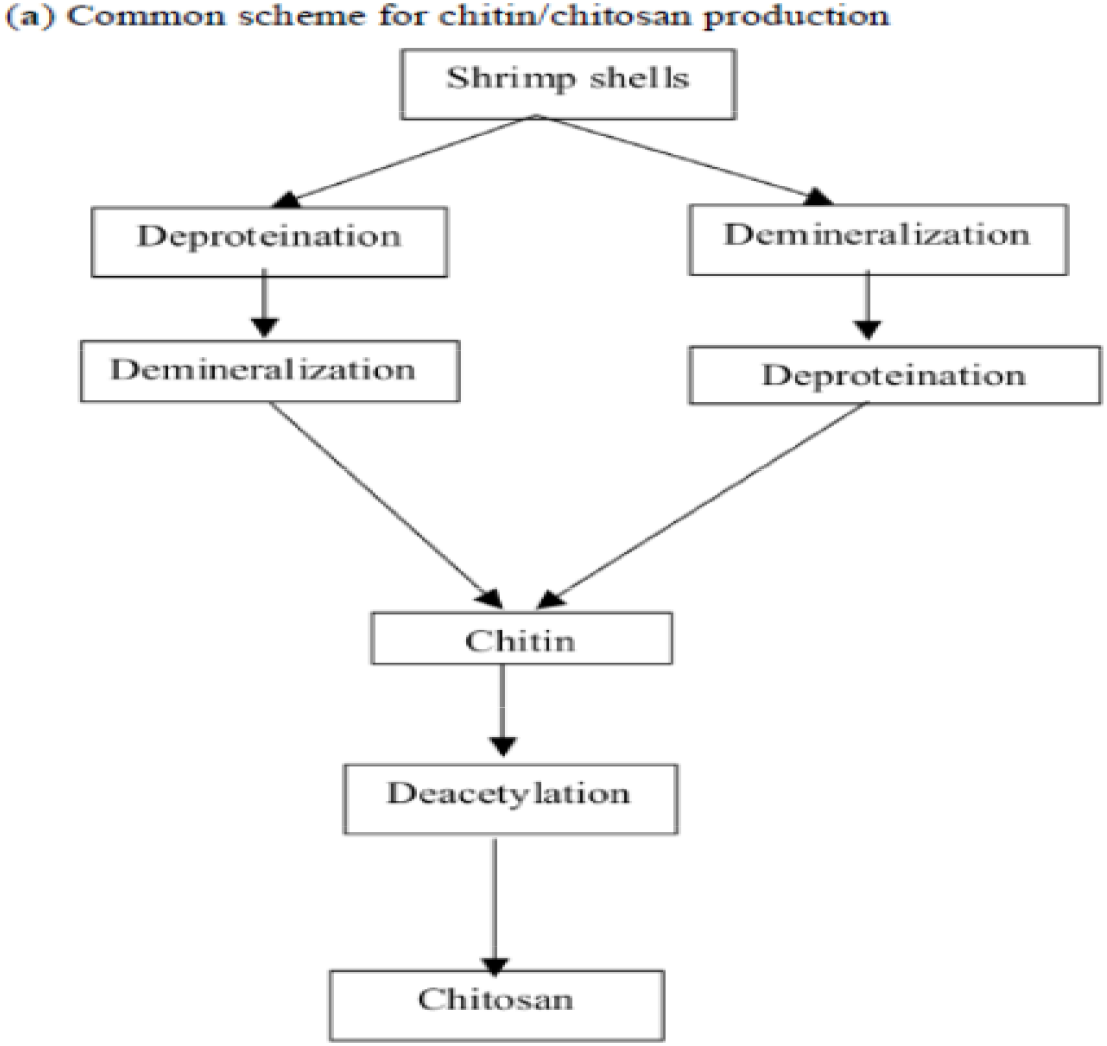
Chitin to Chitosan Process

### Purification of Chitosan

The obtained chitosan has to be purified to make it suitable for the pharmaceutical use. The purification process was designed in three steps namely; removal of insolubles with filteration, reprecipitation of chitiosan with 1 N NaOH, and demetallization of retrieved chitosan

#### Removal of insolubles with filtration

The 1 mg/ml chitosan acetic acid 1% (v/v) solution is prepared by a magnetic stirrer until an homogenous solution is obtained. The insolubles were removed by filteration through Whatman filter paper 22μm.

#### Reprecipitation of chitosan with NaOH

Chitosan was precipitated from filtered chitosan solution by titration with 1 N NaOH until pH value of 8.5. The chitosan obtained is washed several times with distilled water by centrifuging at 8,000 to 10,000 xg. All the above steps were carriedin the presence of reducing agent Dithiothreitol, (DTT) in order to provide more consistency and reproducibil-ity between chitosan batches for biomedical applications (any other hydroxides other than NaOH are reactive which would another step in purification if such materials are used).

#### Demetallisation of retrieved chitosan

Reprecipitation precedes demetallisation by the addition of 1 ml of 10% w/v Aqueous solution of sodium dodecyl sulfate (SDS) and stirring for 30 min for dissolving the protein left over finally. After leaving the solution stirring at room temperature overnight, 3.3 ml of 5% w/v ethylenediaminetetraa-cetic acid (EDTA) was added and stirred at room temperature for 2 additional hours for precipitation of heavy metals with EDTA. The water insoluble chitosan precipitate was collected by centrifugation at 5000xg for30 min using REMI and washed several times with distilled water by resuspending and re-centrifugation for 30 min. the residue obtained is dried in hot air oven at 60 gently to prevent physical damage in the chain structure. The obtained dried chitosan is stored in the dessicator (Qian and Glanville, 2005).

##### Synthesis of Gold Nanoparticles

In a typical experiment, 100 µl of 1.25x10^−1^ M concentrated aqueous solution of chloroauric acid (HAuCl4) will be reduced by heating for 15 min in 100 ml of chitosan solution prepared in 1% acetic acid to yield a ruby-red solution. The ruby red colored solution yielded an absorbance maximum at 520 nm. According to the related study, the higher chitosan concentration (>0.1% w/v) were stable showing no sign on aggregation.

##### Loading of Erythropoietin onto Gold Nanoparticles

The calculated amount of Erythropoietin having 4000 IU/mL will be added to dispersion of gold nanoparticles reduced using 0.2% chitosan (pH of blank gold nanoparticles was 5). The dispersion is incubated for 16 h at 2–8-C followed by ultracentrifugation at 30,000 rpm for 30 min. The pellet thus obtained will be separated from the supernatant solution.

**Figure 3.3.**
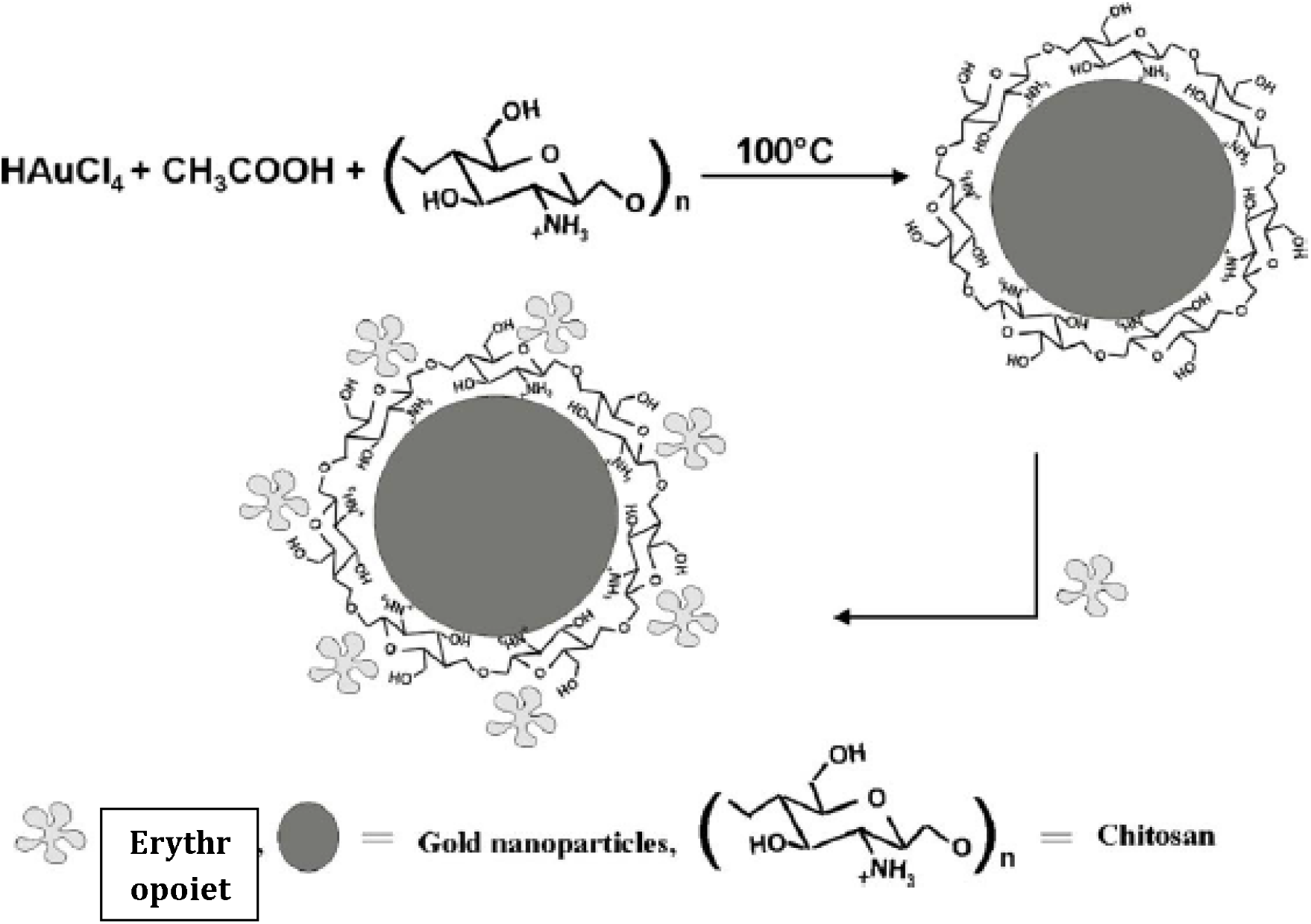
Schematic diagram showing the capping of chitosan on gold nanoparticles and subsequent loading of erythropoeitin on chitosan capped gold nanoparticle

The change in surface Plasmon band of gold nanoparticles, before and after loading of erythropoietin, was monitored by UV-Visible spectroscopy measurements

Calculating the loading efficiency and interpreting the results using statistical analysis

The loading efficiency will be calculated after the free erythropoietin present in the supernatant. Here’s the formula in computing it:

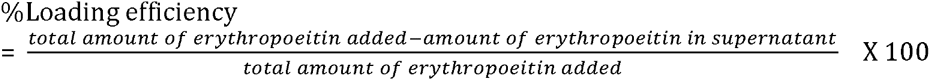

Data will be expressed as mean T S.D. and statistical analysis will be carried out using One-way ANOVA followed by Tukey’s test. A level of significance of P < 0.05 was regarded as statistically significant.

### Converting Chitin to Chitosan

Chitosan, owing to its properties to be biocompatible, absorption enhancer and chemical nature is used as a major part of this study because of its electronegativity that can act as reducing agent to obtain gold nanoparticles. Moreover because of it is a polyelectrolyte it could act as electrostatic stabilizer. It is from converting Chitin from Portunus Pelagicus which has the common name “Alimasag”. It is very popular in the Philippines but just as a delicacy, however it could also be used for nanotechnology. In the process of converting Chitin to Chitosan, it is observed that an average size of Alimasag has 5.5 grams of grinded crab shells that could be used in getting chitosan. There is a yield of 1.9 grams of chitosan from the grinded crab shells where we get 34.55%.

The chitosan undergoes characterization and purification to ensure that it could be used for medical applications then it undergoes the NICOLET 6700 FT-IR Spectroscopy to confirm its functional groups.

**Figure 4.1.**
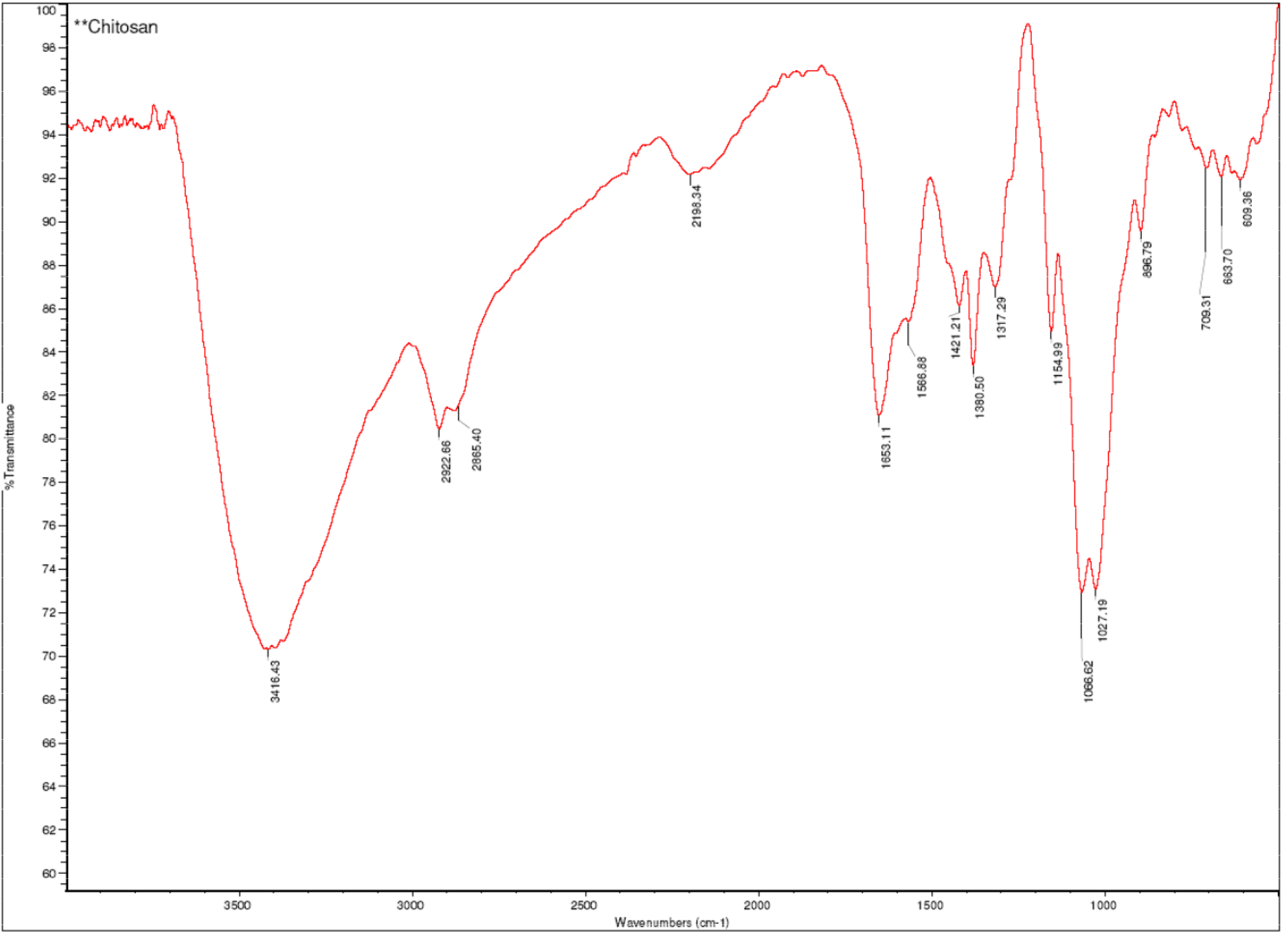
The FT-IR spectroscopy of the obtained chitosan

The FT-IR spectrum of the chitosan showed seven major peaks. The major absorption band is observed between 3400-3250cm^-1^ which represents the C=O stretching in secondary amide. The absorption band between 1250-1020 cm^-1^ shows the free amino group (-NH2) at C2 position of glucosamine, a major group present in chitosan. Further the sample showed the absorption bands for the free amino group between 1027 and 1066 when the peak at 1317 cm^-1^ represents the –C-O stretching of primary alcoholic group (-CH2 - OH). The absorbance bands of 2922 and 2865 indicated the Symmetric CH3 stretching and asymmetric CH2 stretching and CH stretching respectively.

**Table 4.1.** Characterization of Chitosan for Medical applications

| Variables | Standard | Result |
| --- | --- | --- |
| Moisture Content | <10% | 8.69% |
| Ash content | <2% | 1.8% |
| Protein content | <2% | Negative results to Biuret test and Ninhydrin test. |
| Insolubility | <1% | soluble to water and ethanol |
| Viscosity | <5cps | 2cps |
| Heavy metal content | Pb<10ppm | Negative results for Pb |
| Organoleptic properties | No taste, no smell | no smell |

In doing the moisture content, *Total Evaporable Moisture Content of Aggregate by Drying* was used where the original mass of chitosan was 5g then after the process we got a mass of 4.63 grams. Following the formula:

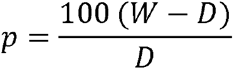

where: p = total evaporable moisture content of sample, percent;

W = mass of original sample, g; and

D = mass of dried sample, g

In the ash content, we get the result of 1.8% which is good and didn’t surpass the 2%. In testing for the presence of proteins, biuret test and ninhydrine tests were used and show negative results. Chitosan was put to water and ethanol to test its solubility and it shows that it is both soluble to ethanol and water. In viscosity, it was put into the pipet to measure how viscous it is, the timer was set and it reached 2 cps. In the presence of heavy metal, lead was the only examined if it wasn’t present to chitosan and it showed negative result which means it was not present in chitosan. In the organoleptic properties, the chitosan had no smell at all.

### Chitosan reduced gold nanoparticles

Gold nanoparticles offer promising gateways in drug delivery. Chitosan reduced gold nanoparticles is regard as potential carrier because of its characteristics. The varying chitosan concentrations (0.1% w/v, 0.25% w/v and 0.5% w/v) are being used to synthesize gold nanoparticles. The concentration of chloroauric acid is 1.48x10^−2^M. The 0.1% w/v shows gel appearance as compare to 0.25% and 0.5% w/v. The 0.25% and 0.5% w/v are more solid in appearance as compare to 0.1% w/v. The 2000 IU/mL of erythropoietin were loaded to the varying chitosan concentrations reduced gold nanoparticles and incubate into 2-8˚C then put into the centrifuge with 13,000rpm for 30 minutes. The pellet thus obtained was separated from the supernatant solution. The volumes of the erythropoietin after the centrifuge were measured.

**Table 4.2.** Varying chitosan concentrations in synthesising reduced gold nanoparticles

|  | Samples |  |  |
| --- | --- | --- | --- |
| Trials (mL) | 0.01% w/v | 0.25% w/v | 0.5% w/v |
| Trial 1 | 0.3 | 0.5 | 0.4 |
| Trial 2 | 0.3 | 0.6 | 0.6 |
| Trial 3 | 0.5 | 0.6 | 0.4 |

After the centrifugation, the varying chitosan concentrations reduced gold nanoparticles result into 0.3-0.5mL, 0.5-0.6mL and 0.4-0.6 mL for 0.1%w/v, 0.25% w/v and 0.5% w/v respectively as what is shown in Table 4.2. The concentrations were done in 3 trials. The highest volume of erythropoietin that could be delivered is 0.25%. It is because of the viscosity as compare to 0.1% w/v and 0.5% w/v. It is tested statistically to know if there’s a significant difference between the chitosan concentrations.

**Table 4.3.** One-way Analysis of Variance of chitosan concentrations

TRIALS 1 to 3
|  | Sum of Squares | Df | Mean Square | F | Sig. |
| --- | --- | --- | --- | --- | --- |
| Between Groups | .060 | 2 | .030 | 3.000 | .125 |
| Within Groups | .060 | 6 | .010 |  |  |
| Total | .120 | 8 |  |  |  |

\*F<4.46

*F<4.46

In One-way ANOVA with 0.05 level of significance shows that there is no significant difference between varying chitosan concentrations. The obtained F ratio did not quite reach the 0.05 level of significance, F(2,8) = 4.46. However, it is noted that 0.25% w/v releases bigger amount of erythropoietin as compare to the other concentrations.

**Figure 4.2.**
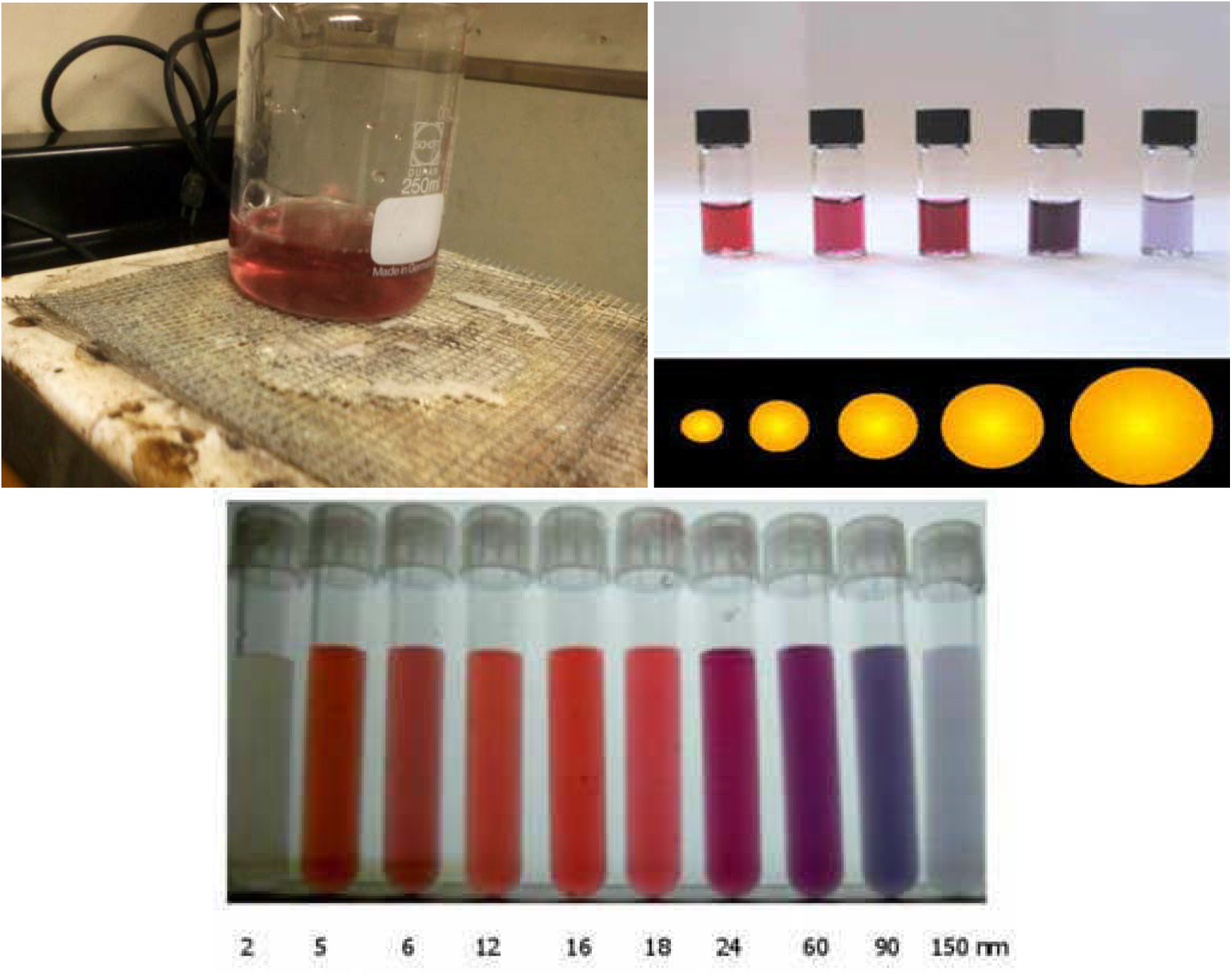
The chitosan reduced gold nanoparticles in comparing to its particle size

**Figure 4.3.**
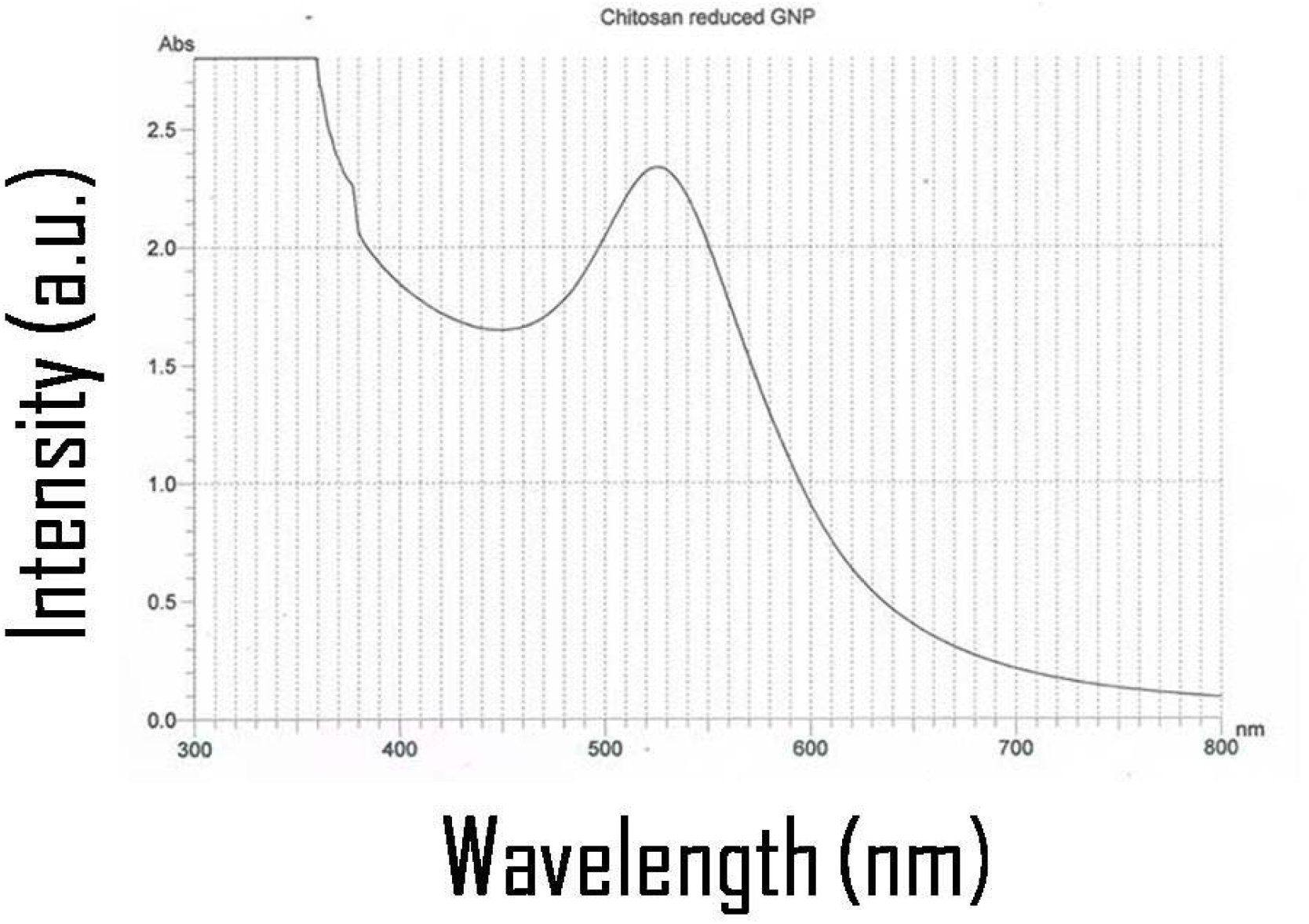
Visible absorption spectra of chitosan reduced gold nanoparticles

In synthesizing gold nanoparticles using chitosan solutions create a change in color which is ruby-red. It pertains to the size of the nanoparticles that were synthesized by chitosan. In Figure 4.2, the chitosan reduced gold nanoparticles give a ruby-red color which shows that the particles obtained are small. The change in color is due to surface Plasmon resonance which is the collective oscillation of electrons in a solid or liquid stimulated by incident light. In the figure, the apex of the visible absorption spectra of chitosan reduced gold nanoparticles show at 526 nm which confirms the presence of gold nanoparticles according to the related researches. Gold nanoparticles exhibits SBP at 526 nm due to collective oscillations of the electron at the surface of the nanoparticles (6s electrons of the conduction band for gold nanoparticles) that is correlated with the electromagnetic field of incoming light, i.e. the excitation of the coherent oscillation of the conduction band.

### Conclusions

In conclusion, the study proves that the chitosan from Portunus Pelagicus is capable of synthesizing reduced gold nanoparticles that could be a potential carrier for the delivery of erythropoietin.

- The obtained chitosan shows seven major peaks (3400-3250cm^-1^, 1250-1020 cm^-1^, 1027, 1066, 1317 cm^-1^, 2922 and 2865 cm ^-1^) which corresponds to its functional groups. It is further characterized to pass the characteristics required for medical applications.
- The varying chitosan concentrations (0.1% w/v, 0.25% w/v and 0.5% w/v) are used in synthesizing gold nanoparticles. Among the concentrations, 0.25% w/v could deliver bigger volume of erythropoietin because of the viscosity as compare to other concentrations. However, there is no significant difference among the concentrations of chitosan reduced gold nanoparticles in delivering erythropoietin.
- The color of the chitosan reduced gold nanoparticles is ruby-red which shows that the size of the gold nanoparticles is small and appropriate for drug delivery.
- There is no change in color before and after the erythropoietin is loaded onto it. The change in color of the chitosan reduced gold nanoparticles is due to surface Plasmon resonance.
- The visible absorption spectra of the chitosan reduced gold nanoparticles shows that gold nanoparticles exhibits SBP at 526 nm due to collective oscillations of the electron at the surface of the nanoparticles. It confirms the presence of the synthesized gold nanoparticles in the solution.

### Recommendation

Based from the discussion of conclusion, the study found some possibilities and imperfection in conducting the study. To refine the study, the researches recommend the following for the enhancement of the study.

- The Alimasag that would be used is better processed immediately. Moreover. It is better to increase the concentration of HCl or NaOH depending on the amount of crab shells that would be processed but not too far from the value given in the procedure.
- In determining the ash content and moisture content, it would be more precise to use oven that has capability of heating in high temperatures. If turbidity is possible to be measured, it could be done because it is a characteristic for medical applications.
- The pellet obtained could be redispersed in milli Q water to further characterize the erythropoietin loaded reduced gold nanoparticles. The free erythropoietin present in supernatant could be determined by ELISA.
- In more precise use of the carrier, it could undergo in vitro diffusion studies for 6 months to observe if there are aggregations in different concentrations.

